# Age-independent influence of hematopoietic stem and progenitor cell populations during hematopoietic reconstitution

**DOI:** 10.1101/2020.08.05.237891

**Authors:** Frauke Gotzhein, Tim Aranyossy, Lars Thielecke, Tanja Sonntag, Vanessa Thaden, Boris Fehse, Ingo Müller, Ingmar Glauche, Kerstin Cornils

**Author notes:** Correspondence: PD Dr. Kerstin Cornils, University Medical Center Hamburg-Eppendorf, Department of Pediatric Hematology and Oncology, Division of Pediatric Stem Cell Transplantation and Immunology and Research Institute Children’s Cancer Center Hamburg, Martinistr. 52, 20251 HAMBURG. FG and TA contributed equally as co-first authors. IG and KC contributed equally as co-senior authors.

## Abstract

Hematopoietic stem cell transplantation (HSCT) following myeloablative conditioning represents the only curative treatment option to overcome numerous hematologic malignancies. Despite the overall clinical success to achieve reconstitution of the blood system in many patients, the length of the aplastic phase until recovery response remains a critical parameter for patient survival. Fatal infections are the major risk factor for immunocompromised individuals and in many cases lead to transplant related morbidity and mortality. Therefore, we hypothesized that transplants from young donors might be beneficial for the spatiotemporal blood reconstitution in comparison to old grafts. Here we report on our multicolor-coded barcode system (BC32) to track hematopoietic stem and progenitor cell (HSPC) populations in a murine model of HSCT. By using different experimental settings of young and aged donor and recipient animals we compared the influence of age on graft composition, leukocyte recovery and more precisely, the contribution of individual HSPC subpopulations to the process. We show that blood reconstitution is substantially driven and maintained by HSCs and MPPs, while the latter accounted for higher output rates than HSCs, confirmed by FACS and NGS analyses. Reconstitution patterns were highly polyclonal and stable in all animals aside from variability between individual animals. This leads to the conclusion that HSPCs home, engraft and differentiate independently of donor and recipient age in our experimental conditions. Our findings add to ongoing efforts to expand the application of HSCT in older individuals while minimizing therapy related risks resulting from intensive myeloablative conditioning or incomplete blood reconstitution.

## INTRODUCTION

Since the first successful hematopoietic stem cell transplantation (HSCT) in 1957 (1) this therapy has been established as standard of care for many (non-) malignant blood diseases and it often represents the only curative treatment available. In the past, a worldwide increase in the demand for HSCTs has been reported, surpassing 50,000 allogeneic transplants annually (2). Since donor age is one of the most predictive, non-HLA donor characteristics for survival after HSCT (3, 4) the frequency of donors with an age below 30 raised up to 69 % in the past years (5). However, multiple studies showed that HSCTs in older patients result in worse outcomes compared to younger patients, particularly due to high rates of transplant-related, non-relapse mortality and morbidity caused by traditional myeloablative regimens or infections (6-8). As a result, elderly patients were excluded from HSCT for long periods of time. Fortunately, with the emergence of less toxic, particularly reduced-intensity conditioning regimens (RICs), the application of HSCTs as a potential treatment option for increasingly older patients has been reevaluated over the last decades (9, 10). This raises the question how age of donor and recipient affect HSCT success on a cellular level and if this process might be accelerated in order to alleviate infections in the early post-transplant phase.

Besides the influence of donor and recipient age also the different cell sources seem to impact the success of HSCTs. There are currently three different cell sources available, namely bone marrow (BM), mobilized peripheral blood stem cells (PBSC), and umbilical cord blood (UCB). Depending on the patient and the setting (autologous or allogeneic transplantation) the majority of HSCTs is conducted with BM or PBSCs (11). Unfortunately, due to a lack of HSC-specific markers, the true number of HSCs in a graft is still not measurable. If medically indicated, the positive identification of the cell-surface marker CD34 is used to enrich for HSCs. However, the CD34 protein is not exclusively expressed on HSCs, but also on a variety of different progenitor cells (12). Therefore, a CD34^+^ cell population is extremely heterogeneous, most likely comprised of a large variety of different hematopoietic progenitors and only a small fraction of bona fide HSCs (13, 14). Even though it is possible to further differentiate the particular subpopulations utilizing additional phenotypic marker combinations, the exact composition of a clinically applied human stem cell graft still remains unknown. Therefore, it remains elusive how different subpopulations of the graft influence hematopoietic reestablishment and how the composition of subpopulations could be affected by the donor and recipient age.

In recent years, the development of genetic barcodes as a means of cellular labeling has revolutionized clonal tracking (15, 16), especially in the field of hematopoiesis (17). Using integrating retroviral vectors, artificial genetic sequences (termed barcodes) can be introduced in the genome of certain cells of interest as a cell-specific and permanent marker. For instance, HSPCs can be labeled prior to HSCT, and since those genetic labels are inheritable, the productivity of the initially marked cells during and after reconstitution can be measured by the abundance of the respective genetic barcode within the hematopoietic “organs” (e.g. bone marrow, spleen) as well as in sorted mature blood subpopulations.

The majority of the published work on clonal barcoding utilized highly purified murine HSC-populations, sorted according to respective marker profiles (18-21). Other longitudinal studies of blood reconstitution, working with humanized mouse or primate models, investigated grafts consisting of a mixed CD34^+^ cell population, similarly to a standard human transplantation setting (22-30). However, the difference in clonal composition and clonal contribution between highly purified, aged and young HSCs was so far only studied by Verovskaya et al. in a murine model utilizing competitive transplantations (18).

We have recently developed a novel approach to inheritably mark individual cells, which combines the virtues of two formerly established marking techniques, namely, RGB marking (31) and the aforementioned genetic barcoding. While the fluorescence-based RGB marking allows for a phenotypic distinction of different cell clones *in situ*, the genetic marking facilitates the robust and long-term follow-up even if the expression of the fluorescence genes is diminished or extinct. This combination of techniques enables a simultaneous analysis of four different cell populations with theoretically up to 4^32^ uniquely labeled clones each (32, 33). In the past, we already successfully utilized this approach to investigate different viral vectors and the influence of the respective promoter on the reconstitution of the peripheral blood (34).

In the study presented here, we examined to which extend different hematopoietic subpopulations drive the reconstitution after HSCT on a clonal level. To this end, we applied our marking approach to individually label four different BM subpopulations and to follow their distinct clonal output during reconstitution. We particularly focused on the contribution of cells from young and aged donor mice transplanted into young and aged recipients. To obtain results that are more comparable to the human situation, we used similar combinations of surface markers for the labeling of corresponding subpopulations in a human graft. Our experimental data provide insights into the clonal output of the initially transduced cell populations to better understand the temporal reconstitution processes after HSCT.

## METHODS

### Generation and production of barcoded lentiviral vectors

Based on the original LeGO vector constructs, we generated lentiviral vectors encoding for eGFP and three derivatives, namely eBFP, Venus and T-Sapphire as described in Aranyossy et al. (34). All vector constructs were additionally equipped with the BC32-barcode sequences (barcoded plasmid library), to add a genetic label and to also cipher the fluorescent protein in the BC32-backbone (Figure 1A). VSV-g pseudotyped lentiviral particles were produced as described, concentrated via centrifugation and titrated on HEK293T cells (35).

**Figure 1:**
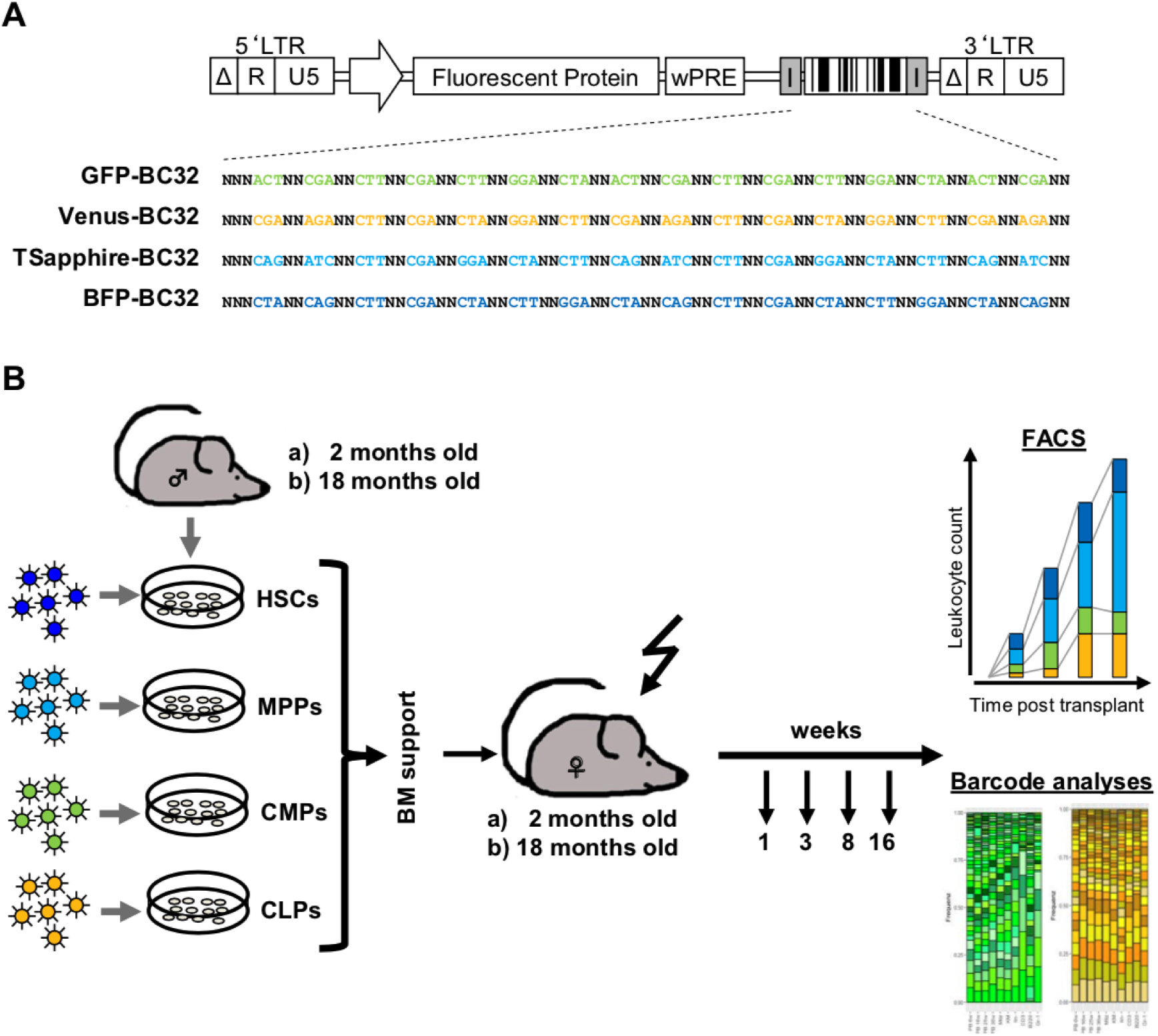
Vector construction and experimental setup. (A) Four different barcoded plasmid libraries based on lentiviral vectors encoding for the fluorescent proteins GFP, Venus, T-Sapphire and BFP were used to produce viral supernatant. The color of the fluorescent protein is also ciphered by the order of the fixed triplets (34) (Aranyossy 2017). (B) For the murine transplantation experiments, lineage-negative cells from male donor mice (n=30; 2 months or 18 months old), were isolated from bone marrow and sorted by FACS into hematopoietic stem cells (HSCs), as well as the following progenitors: multipotent progenitors (MPPs), common myeloid progenitors (CMPs) and common lymphoid progenitors (CLPs). The four different populations were transduced independently with the 4 different lentiviral constructs depicted in (A). After short-time transduction the cells were mixed and transplanted together with bone marrow support into lethally irradiated female recipients (n=30; 2 or 18 months old). 1, 3, 8 weeks post transplantation and at final analysis (16 weeks post transplantation), a small group of animals (n≤7) were sacrificed and the hematopoietic organs were taken for flow cytometry analysis and DNA extraction for barcode retrieval.

### Isolation and transduction of target cells

All animal procedures were conducted in accordance with the regulatory guidelines and were approved by the local authorities (Behörde für Gesundheit und Verbraucherschutz-Veterinärwesen/Lebensmittelsicherheit) and the University Medical Center Hamburg-Eppendorf (UKE). Food and water were available *ad libitum*.

Thirty male C57Bl/6 mice were used as donors. Young donor mice were two months old, while old donor mice were 18 months old. BM of the donors was harvested by crushing femur, tibia and ilium. Lineage negative cells were isolated using Miltenyi’s lineage depletion kit according to manufacturer’s instructions. After staining with an established antibody panel, cells were sorted using BD FACS AriaIllu (BD Biosciences) into hematopoietic stem cells (HSCs), multipotent progenitors (MPPs), common myeloid progenitors (CMPs) and common lymphoid progenitors (CLPs) (Supplementary Data 2), and subsequently transduced with the respective barcoded vector via spin-infection in StemSpan medium containing 50µg/µL Stem Cell Factor (SCF, PeproTech). Transduction efficiency was measured via flow cytometry and quantified via expression of the respective fluorescent protein three days post transduction. Alternatively, we fitted a Poisson distribution to barcode numbers from single cell cultures to provide an estimate of the overall transduction efficiency.

### Murine transplantation experiment

After one day of transduction, the four cell populations were washed and merged. Together with BM support (1 x 10^6^ cells of female donors), the cells were transplanted into the recipient animals via tail vein injection. Thirty female mice were taken as recipients and lethally irradiated prior to transplantation (9.5 Gy). To follow the course of reconstitution, groups of five to seven animals were taken out of the experiment at one, three, eight and 16 weeks post transplantation. Single cell suspensions from peripheral blood (PB), BM and spleen were isolated using QIAamp DNA Blood Mini Kit (Qiagen) and analyzed by flow cytometry BD LSRFortessa (BD Biosciences) for fluorescent protein expression. Cells from spleen were additionally stained for CD3, B220, CD11b and Ly6G to determine the content of marked cells in the T cell, B cell, granulocyte and monocyte/macrophage compartment (Supplementary Table 1). Cells were also used for DNA extraction from whole spleen (see above, Figure 1B).

Single cells derived from HSCs or MPPs were sorted according to their fluorescent protein expression (compare Figure 2B). Up to 14 cells were sorted per FP color, and the barcode sequences were PCR-amplified directly from the single cells in a nested PCR approach. To estimate the number of barcodes per cell, we analyzed the Sanger sequencing chromatogram (36).

**Figure 2:**
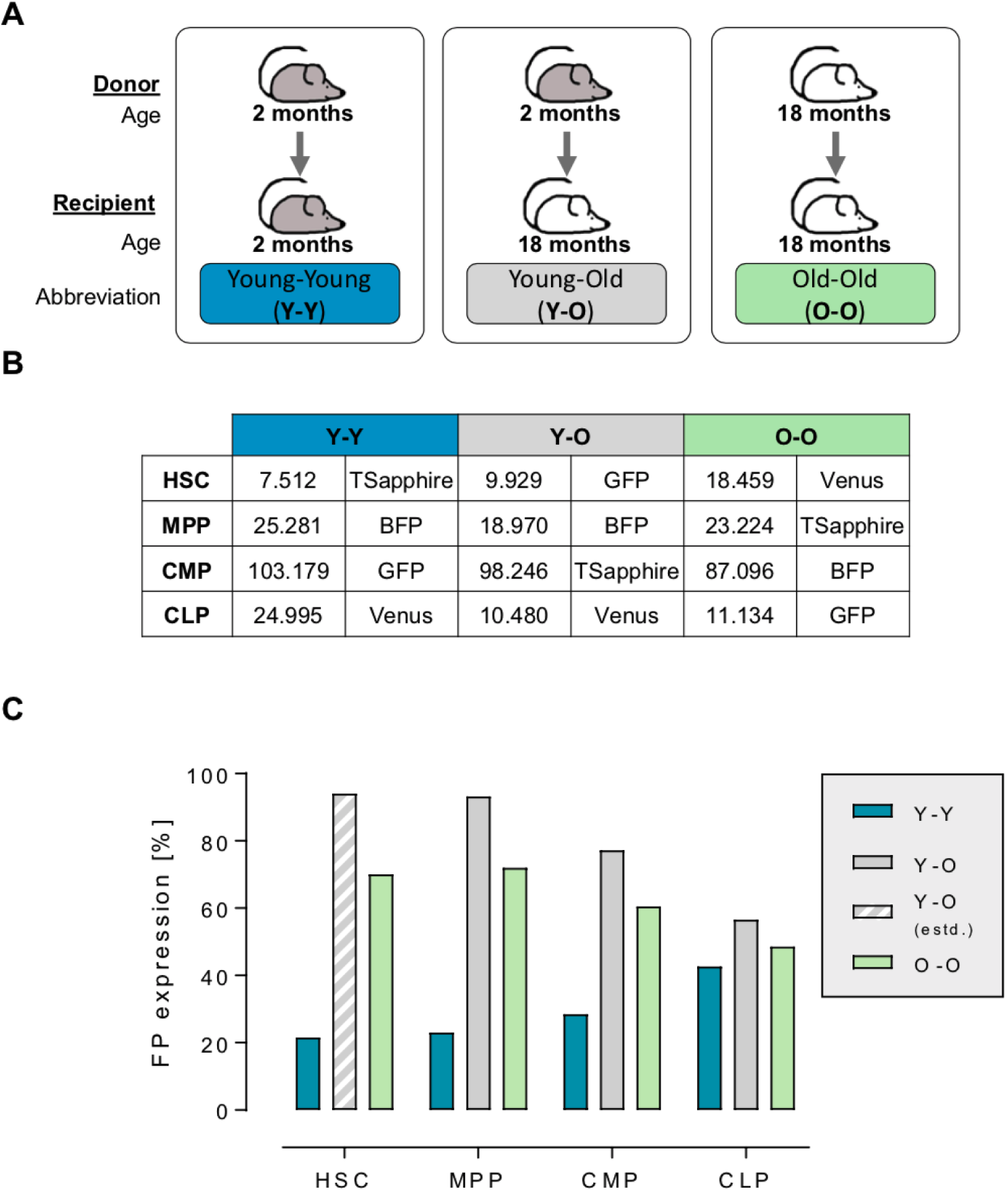
Experimental groups, sorted cell numbers and transduction efficiencies. (A) The experimental groups were set up as follows: cells from 2-month old male mice were used for transplantation into 2-month old female recipients (blue: Y-Y), cells taken from old donors (18-month old) were transplanted into young recipients (2-month old, grey: Y-O) and finally cells from old donors were transplanted into old (both 18-month old) recipients (light green: O-O). (B) According to the marker profile, cells were separated by FACS-sorting and transduced with lentiviral barcode-vectors encoding for the depicted fluorescent protein. Numbers depict the mean amount of cells obtained for each subpopulation. (C) Three days post transduction, transduction efficiencies for the four subpopulations were measured by flow cytometry of the respective fluorescent protein. The value for the transduced HSCs in the Y-O experiment was estimated (grey striped bar). Blue bars represent Y-Y experimental setting (n=3-5), grey bars represent Y-O experimental setting (n=4-7) and light green bars represent O-O experimental setting (n=4-6).

### Determination of chimerism

200 ng of genomic DNA from BM of the transplanted animals was used to determine the chimerism. We performed droplet digital polymerase chain reaction (ddPCR) in a duplex reaction by simultaneously amplifying a Y-chromosome-specific fragment and a control amplicon (erythropoietin receptor). Analyses were performed using the QX100 system of BioRad according to manufacturer’s instructions (32, 37).

### Barcode analyses

The genetic barcodes were extracted and amplified via PCR from the genomic DNA as described earlier (33, 34). After demultiplexing, the NGS results were analyzed utilizing an in-house developed R-package (33, 38). The detected sequences were first sorted according to their backbone sequence, thereby allowing for a distinction of the different transduced cell populations. Then, single reads were quality checked, barcode abundancies were calculated and an error-correction was applied to cope with PCR- and NGS-introduced errors. Subsequent measurements and clonal tracing within the same recipient were not possible due to the limited amount of extractable material. Therefore, we focused our analyses on the cell type-specific number of clones contributing to regenerate hematopoiesis.

## RESULTS

### Graft production

We established a graft, in which populations of HSCs, MPPs, CMPs and CLPs were distinguishable based on the virally integrated fluorescence marker and the barcode backbone. Furthermore, by analyzing the actual genetic barcodes we were able to additionally identify individual clones within each of those four populations.

In order to test the influence of different conditions regarding donor or recipient age we outlined three different experimental transplantation settings, namely cell populations harvested from a young donor transplanted into young recipients (Y-Y), young donor cell populations transplanted into old recipients (Y-O) and old donor cells transplanted into old recipients (O-O) (Figure 2A). Technically, the BM of 30 young and old male donor mice was used for the enrichment of lineage negative cells followed by subsequent fluorescence-activated cell sorting (FACS) to further separate HSCs, MPPs, CMPs and CLPs (compare Figure 1B). While we obtained similar cell numbers between young and old donors for the MPPs and CMPs, the numbers for the HSCs were almost two times higher when derived from the aged donor mice (Figure 2B). Additionally, the CLP amount in the Y-Y setting was more than two times higher compared to both other experimental settings.

Next, sorted cell populations were transduced individually with one of the four barcoded lentiviral vector libraries. In order to minimize potential backbone-related systematic bias, we changed the different fluorescence proteins between the experimental settings (Figure 2B). We aimed to achieve a transduction rate of around 20% to ensure single vector copy numbers (39). While in the Y-Y experimental part, we measured moderate transduction rates of 20 – 40%, the transduction efficiencies in the other two parts exceeded the target range, reaching levels of about 50% (Figure 2C), for which multiple integrations per cell become more likely. We determined the average vector copy number (VCN) in HSC- and MPP-derived cells from those two experiments (O-O and Y-O) using a sequencing chromatogram (36). For the O-O setting we obtained a corresponding estimate of VCN=2 (HSC and MPP) and a VCN=3 for the Y-O setting (only MPP) (Supplementary Data 1). For the Y-O HSCs, we used the VCN from the single cells and applied a Poisson statistics to estimate the transduction efficiency as we could not directly measure the VCN via flow cytometry due to low cell numbers (Figure 2C).

### Dynamics of blood reconstitution from old and young grafts

After the transplantation, we first confirmed that the engrafted cells were of male origin and therefore derived from the transplanted graft. To this end, we performed ddPCR on the Y-chromosome to determine male/female chimerism in the BM after transplantation. Although we observed great inter-individual variances, the steady increase of male cells in the recipient mice was comparable between the three groups (Figure 3A). On average, we detected 68% of the transplanted male cells in the bone marrow of the recipients in the Y-Y experiment, 44% in the Y-O and 56% in the O-O experiment after 16 weeks (Figure 3A). Next, we analyzed the spleens at each time point via flow cytometry to monitor the overall reconstitution of different blood cell subpopulations, namely T cells (stained by CD3e) and B cells (stained by B220) from the lymphoid compartment and monocytes/macrophages (stained by CD11b) and granulocytes (stained by CD11b and Ly6G) from the myeloid compartment. As shown in Figure 3B, total leukocyte counts increased until three weeks post transplantation and then stabilized throughout the analyzed experimental time period. No significant differences were observed in the reconstitution dynamics, indicating independence of the age of the donor material (Figure 3C). B cells expanded rapidly and constantly, whereas the other subpopulations stayed roughly stable during the observation time of 16 weeks.

**Figure 3:**
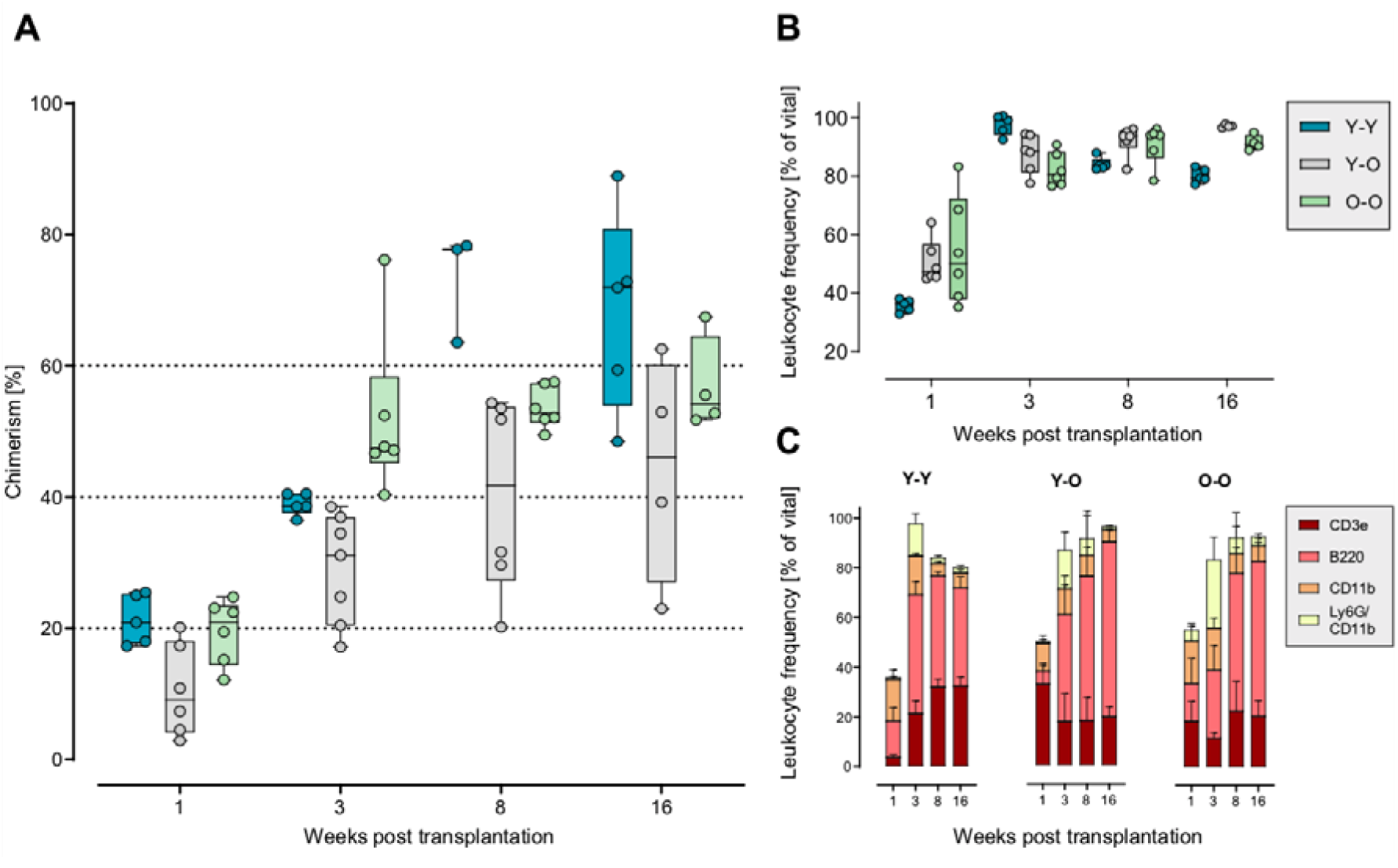
Reconstitution dynamics. (A) We determined the chimerism via a Y-chromosome specific ddPCR in the bone marrow of transplanted animals. Despite great inter-animal variances, we did not observe significant differences in the reconstitution kinetics. (B) Total leucocyte counts were measured from spleens. (C) Single-cell suspensions from spleen were used to stain for hematopoietic subpopulations, like T cells (CD3e), B cells (B220), monocytes/ macrophages (CD11b) and granulocytes (CD11b and Ly6G). Frequencies of these populations after transplantation were similar, independent of the age of donor mice (Y-Y and Y-O vs. O-O). Blue bars represent Y-Y experimental setting (n=3-5), grey bars represent Y-O experimental setting (n=4-7) and light green bars represent O-O experimental setting (n=4-6).

### Contribution of marked cell populations to the reconstitution

In order to determine how overall hematopoietic reconstitution was driven by the initially marked stem and progenitor cells, we assessed the abundance of their progeny in the spleen by gating on the expression of the respective fluorescent protein (Figure 4). In the Y-Y experimental setting, overall contribution was found to be extremely low, as FP expression values mostly ranged below 2%. An exception was week three, in which we detected an anomalous high contribution of MPP- and CLP-derived cells in all mice. Sixteen weeks after transplantation, the expression of fluorescence markers declined to nearly zero (Figure 4A). On the contrary, in old recipients (Figure 4B-C), we detected higher amounts of marked cells. In both groups, the contribution of HSCs and MPPs peaked at around 10% at the later time points. Within the Y-O setting, we observed an almost stable MPP contribution during the entire observation time. This was not present in the O-O experiment, where the MPP-derived progeny declined after a peak at week three. In general, compared to HSC-derived cells, MPP output was consistently higher at all time points, independent of donor or recipient age. After the first three weeks, we observed differences between the Y-O and O-O group: in the young grafts, the peak of HSC progeny appeared eight weeks post transplantation, whereas in the old grafts we observed an earlier peak three weeks post transplantation. The contribution of CLP-derived cells was barely detectable at all time points, while a distinct contribution of CMPs was detectable at week one post transplantation in both experimental groups, which diminished at three weeks. Notably, in the Y-O group, we observed a high variability between animals compared to the O-O group, especially at later time points. A prominent example is the amount of MPP-contributing cells at 16 weeks (Y-O: 7.9 ± 7.67 in comparison to O-O 3.0 ± 1.03).

**Figure 4:**
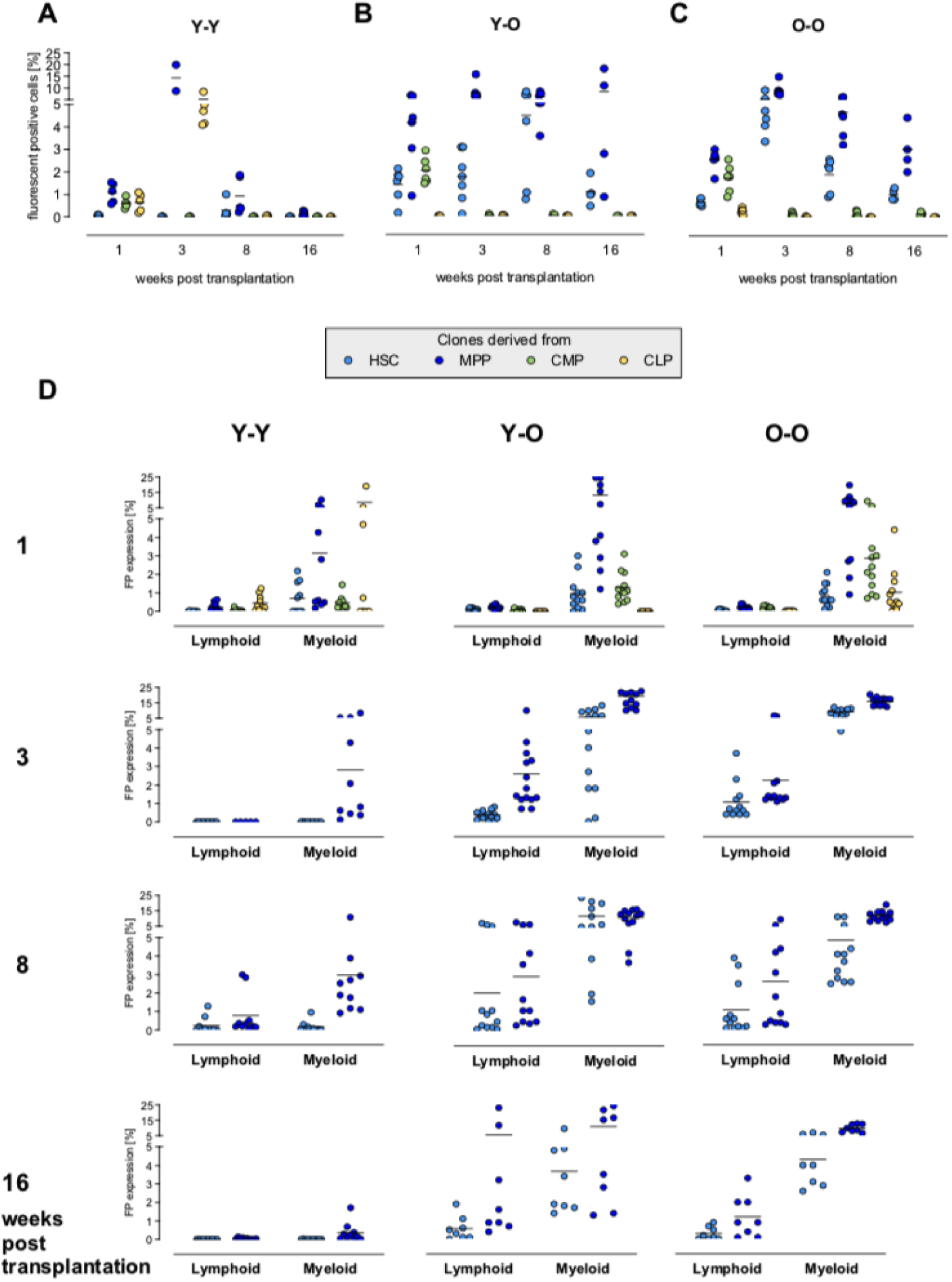
Contribution of marked cell populations. (A, B, C) Single cell suspensions from spleen were used to determine the contribution of the initially marked cell populations via their fluorescent protein expression. The contribution of HSC-, MPP-, CMP- or CLP-derived cells in the peripheral blood was measured according to the respective FP expressed. (D) FP expression was measured in the lymphoid (T cell (CD3) and B cells (B220)) and in the myeloid (monocyte/macrophage (CD11b) and the granulocyte (CD11b/Ly6G)) compartment, also by FACS. The spleen was mostly comprised of HSC- and MPP-derived cells. CMP-derived cells were solely detectable 1 week after transplantation. CLP-derived cells were barely detectable. No data was available for CMP- and CLP-derived cells 3, 8 or 16 weeks post transplantation due to low/no cell numbers. Light-blue dots: HSC-derived clones, dark-blue dots: MPP-derived clones, green dots: CMP-derived clones and yellow dots: CLP-derived clones.

Next, we analyzed the contribution of the differently marked cell populations into distinct downstream compartments, namely the lymphoid (as assessed by their contribution to mature T and B cells) and the myeloid (granulocytes and monocytes/macrophages) in all spleen samples (Figure 4D). In line with the previous results, the main contribution detected in all mature blood cell lineages was largely driven by the marked HSCs and MPPs, starting to become evident already one week after transplantation. Also, at week one we detected CLP-derived cells in the lymphoid compartment (Y-Y), but surprisingly also in the myeloid compartment. The latter was only evident in the Y-Y and in the O-O experimental groups. In the Y-O group, we could not detect any CLP-derived cells at all. CMP-derived cells were solely contributing to the myeloid compartment at week one post transplantation, and vanished with increasing experimental observation time.

We could not detect any long-term contribution of CLPs and CMPs, leading to the conclusion that these cells were only present intermittently and did not contribute to the engrafted subfraction of cells (Figure 4D; 3-16 weeks post transplantation).

The progeny of the marked MPPs and HSCs contributed to both myeloid and lymphoid cells in the spleen of all recipient animals and in all three groups. For the Y-Y experiment, we could detect both, MPP- and HSC-output at eight weeks post transplantation in the lymphoid as well as in the myeloid compartment, whereas at all other time points, the contribution to the lymphoid cell population was barely detectable. In the myeloid compartment, we detected reliable numbers of MPP-derivatives over the whole observation period.

In both experiments with old recipients, MPPs and HSCs contributed to all observed cell populations. With regard to the age of the donor cells, we did not observe evident differences in the contribution of marked cells after transplantation. MPP-derived cells were detectable at all time points, with a higher contribution to the respective compartment (myeloid or lymphoid) in comparison to the HSCs. The only remarkable difference between the groups with old recipients is the variability between the animals. This is primarily seen at the last time point, 16 weeks post transplantation, as already described earlier. Whereas the values for the FP expressing cells in the O-O experiment group together, especially in the MPP-derived myeloid cells (Figure 4D, lower right plot), the inter-animal and cell type specific variances in the Y-O experiment are distinctly higher (MPP in Y-O: lymphoid 5.3 ± 8.02 and myeloid 10.8 ± 9.52 in comparison to O-O: lymphoid 1.2 ± 1.13 and myeloid 9.6 ± 2.48, Figure 4D, middle plots). Taken together these data indicate, that with comparable experimental conditions mature blood cells reconstitute mostly independent of donor and recipient age with a higher contribution of MPPs over HSC-derived output. As expected, CMP- and CLP-derived output was only detectable in short-term reconstitution, confirming the incapability of the progenitor cells to contribute to long-term reconstitution.

### Clonal composition of reconstituted compartments

To assess the amount of clones driving reconstitution, we analyzed the clonal composition in PB, BM and spleen. Based on the different barcode backbones, we identified the different cell populations and counted the unique clones derived from the initially marked cells. Due to the experimental setting, we could not follow individual clones over time, but used resulting barcode abundances as a surrogate for clonality at any given time point. In Figure 5, the mean number of barcodes per animal is depicted, arranged by organ and precursor population. For a better interpretability of the different kinetics, we linked the independent data points. Although the overall recovery of marked cells was influenced by different transduction efficiencies, we observed a polyclonal reconstitution in all of the three groups. Despite some variability between organs, the overall picture of clonal kinetics was similar. Most HSC-clones were detectable up to three weeks post transplantation, with diminishing numbers over time. We observed the same aforementioned pattern for MPPs, the numbers of contributing clones were higher in comparison to the HSCs and appeared to be more stable. The majority of CMP-derived clones were only present in the first week, with merely a few clones detectable in the later samples. A similar picture was seen in the CLP population, only a few clones were detectable at later time points for the Y-Y and Y-O experimental groups but for the O-O group also a quite remarkable number of clones were visible in BM and spleen at the end of the experiment.

**Figure 5:**
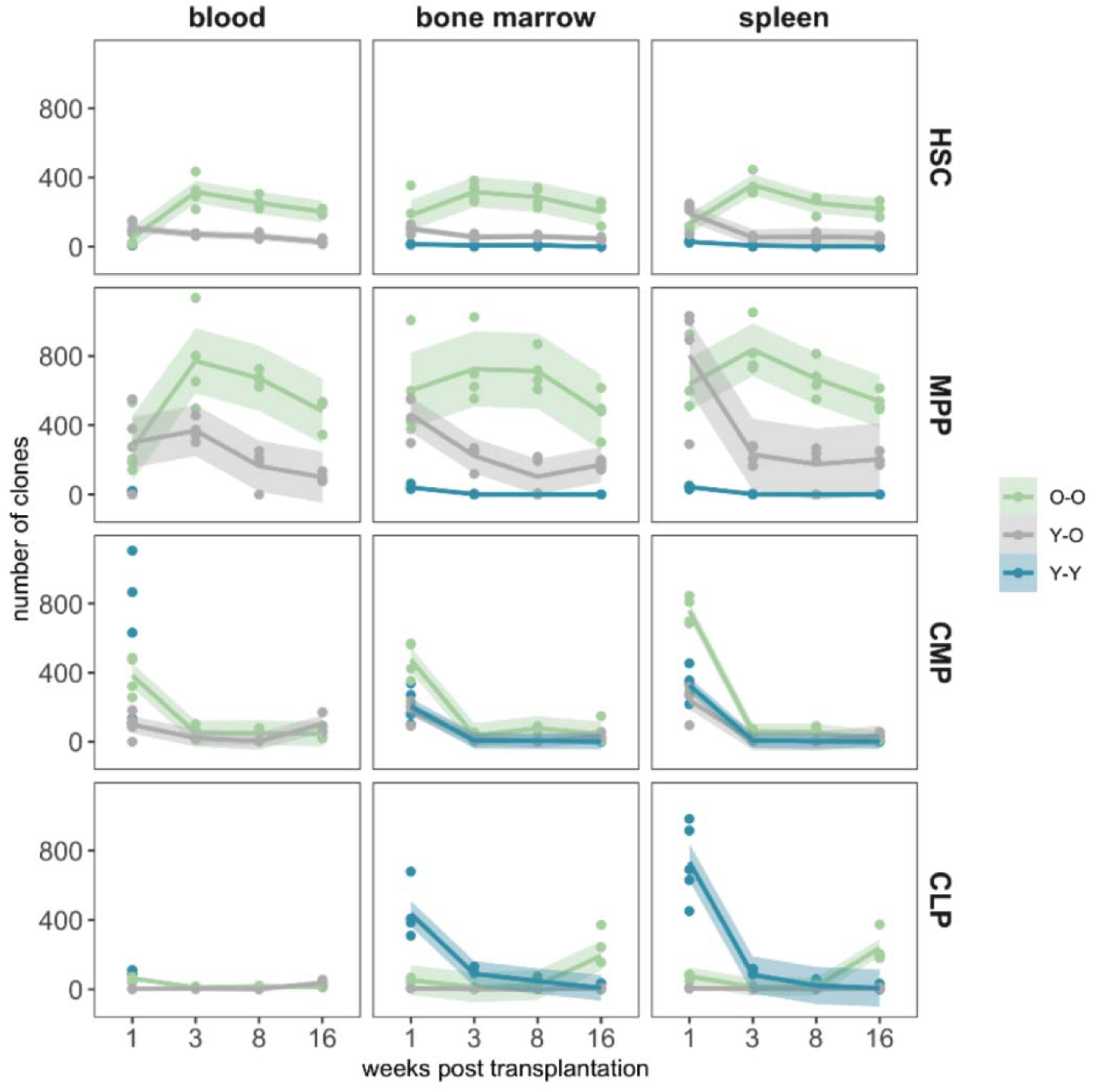
Mean Barcode content. We extracted DNA from peripheral blood, bone marrow and spleen and amplified the BC32 sequences by PCR. NGS and subsequent bioinformatic analyses showed the number of unique barcodes from the initially marked cell population in the respective organ. For a better representation of the reconstitution dynamics, we overlayed the individual clone numbers (points) with fitted local polynomial regression curves (lines) including their respective 95% confidence bands (shaded regions).

In a direct comparison of the reconstitution dynamics of FP-expressing cells (Figure 4A-C) and the number of contributing clones (Figure 5) in the spleen, we observed several similar patterns while others differed strongly. For the HSCs, we detected an increase in FP-expressing cells in a few animals in the Y-O setting at week eight (Figure 4B), while the number of unique clones remained stable. Within the Y-Y or O-O experimental parts, the kinetics of HSC-deviates in terms of FP-expressing and number of clones were similar for each group. Whereas the FP-expression in MPP-derived cells remained almost stable in the Y-O group, the number of barcode-labeled clones declined over time. In the Y-Y experiment, we observed a peak of FP-expressing cells three weeks post transplantation in two animals, whereas the number of barcodes was already comparable between all animals. In contrast to the analyses by flow cytometry, we detected a clonal contribution of CMP- and CLP progenitors to the later reconstitution (after three weeks) with the more sensitive genetic barcode system.

## DISCUSSION

In this study we used our multicolor barcode system to investigate the contribution of four distinguishable HSPC subpopulations, namely HSCs, MPPs, CMPs and CLPs, that were combined in one graft (34). In particular, we choose three different settings of transplantation (Figure 2A) to investigate whether donor or recipient age affects the overall engraftment and the temporal contribution of these subpopulations. We observed that the number of HSCs sorted from bone marrow of old donors was higher in comparison to the young donor material (Figure 2B) (40-42), but following the resulting chimerism and reconstitution dynamics of the (total) leukocytes, we did not detect considerable differences (Figure 3).

As expected, CMP- and CLP-derived cell populations only contributed to short-term reconstitution. They were mainly detectable in the first days post transplantation, probably resulting from a short life span or limited engraftment capabilities of these progenitor cells. Since we did not analyze the presence of CLP-derived clones in the thymus, the observed occurrence of CLP-derivatives at 16 weeks might be explained by the presence of mature T cells. The main contribution to all blood cell lineages was maintained by the HSCs and MPPs (Figure 4), independent of the donor or recipient age. Although we were confined to a co-transplantation setting, our data indicates that cells from the MPP pool can possess long-term reconstitution potential, and therefore can be able to substantiate HSC functionality.

Our results are in agreement with other findings on MPP contribution. By transplanting different defined subsets of MPPs, Oguro et al. and Lui et al. found that not only HSCs, but also MPPs represent a heterogeneous cell pool that is capable of transient and long-term reconstitution (43, 44). Based on extensive immunophenotyping studies of Wilson et al. (45) and further characterization of proteome, transcriptome and methylome data (46), MPPs and their subclasses (MPP1-MPP4) were thoroughly described and functionally characterized. In our case, the MPP population was sorted based on CD150 expression, thereby resembling the MPP1 and MPP2 subclasses and providing a less lineage-restricted/differentiated form of MPPs. In our experiments, MPP-derived cells showed a higher prevalence during the reconstitution phase, both by FP expression (Figure 4) and in terms of barcode-based clonality analyses (Figure 5). Of note, more MPPs were initially present in the younger grafts (experimental group Y-Y and Y-O). The higher number of MPP-derived clones in the O-O experiment can be explained by an up to three-fold reduced activity of the respective HSCs, as described earlier (47, 48).

Further studying the clonality of the reconstruction process within the four subpopulations, we found a stable polyclonal pattern. Even though our experimental setup did not allow to obtain time course measurements within individual animals, the overall clone numbers did not support the idea of only few contributing cell clones or a rapid monoclonal conversion.

In summary, within our murine transplantation experiments, we could not detect any profound impact of donor or recipient age during the reconstitution phase over 16 weeks, besides the known reduced activity of aged HSCs. It is a challenging question whether those results depict the clinical situation. A direct comparison of our results with patient data faces different obstacles, especially since data on the clonality of human blood reconstitution are only available from gene therapy studies, usually involving younger patients, in which CD34^+^ HSPCs are gene modified by integrating viral vectors (49, 50). However, it would be of great interest to analyze the susceptibility to lentiviral transduction for different HSPC populations among the heterogeneous CD34^+^ cell pool to elucidate the impact of genetically modified subpopulations on the success of transplantation in general and gene therapy in particular. Until now, the composition of the CD34^+^ cell population in patient’s graft has not been further examined with regard to the frequency of bona fide HSCs and MPPs. Our results indicate, that both populations play an important role for sustained multi-lineage engraftment, whereas donor or recipient age might be less of a limiting factor.

## ACKNOWLEDGEMENTS

First of all, we thank the animal facility of the University Medical Center Hamburg-Eppendorf (UKE) for their support during the time for aging of the mice as well as the experimental procedures. Furthermore, we thank the UKE FACS Core Facility for technical support. Additionally, we want to thank Dr. Andreas Dahl and the Deep Sequencing Group of the BioTec / CRTD at the TU Dresden for conducting the Illumina sequencing.

## FUNDING

This work was supported by the Dr. Werner Jackstädt-Stiftung and the Deutsche Forschungsgemeinschaft (DFG) granted to KC and IG [DFG: CO 1692/1-1 to KC and GL 721/1-1 to IG).

## Notes

### Competing Interest Statement

The authors have declared no competing interest.

