## Supplementary Material for "Age-independent influence of hematopoietic stem and progenitor cell populations during hematopoietic reconstitution"

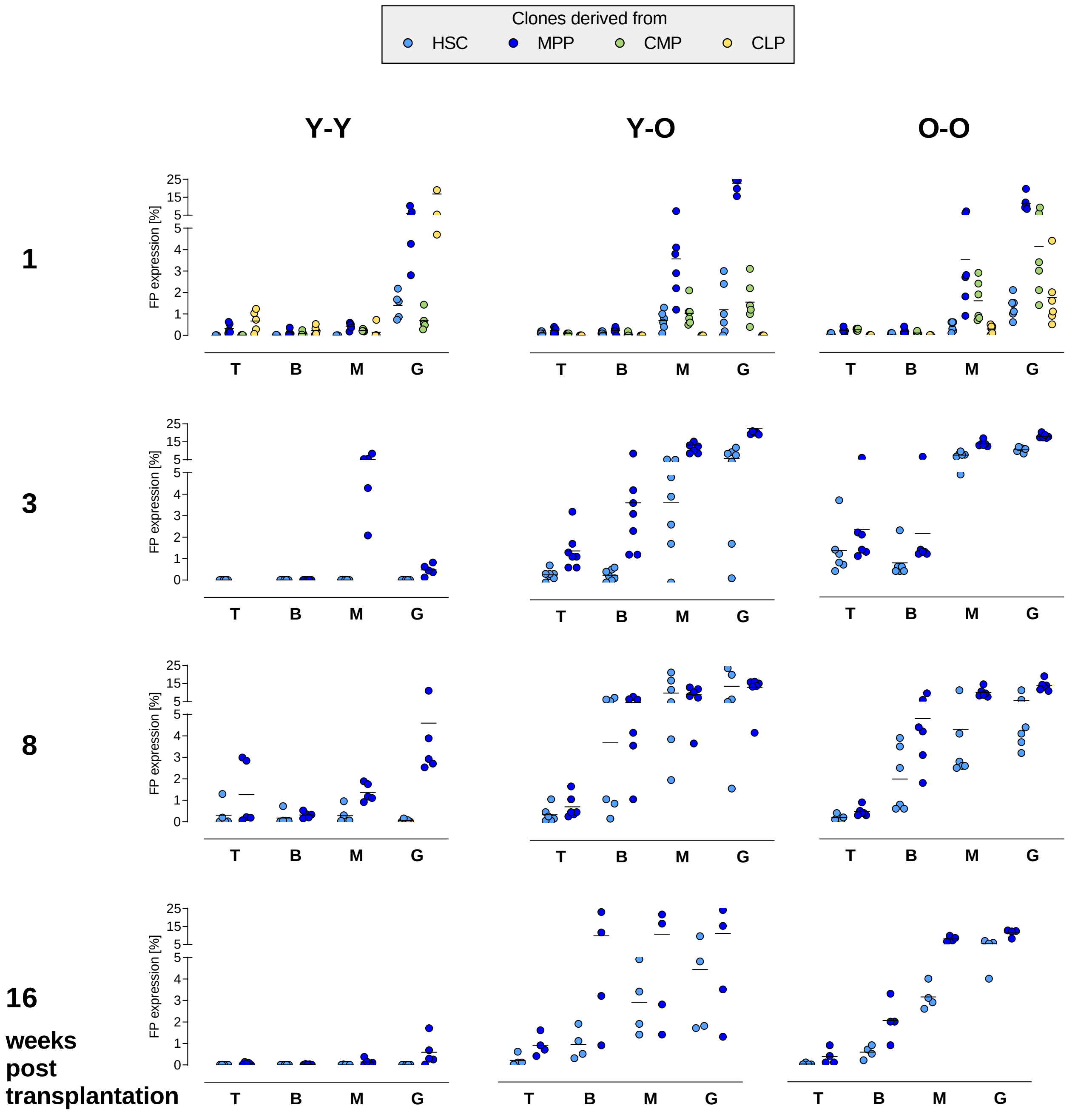

**Supplementary Figure 1: Contribution of marked cell populations in mature blood cell compartments**
FP expression was measured in the T cell (T; CD3), B cell (B; B220), Monocyte/Macrophage (M; CD11b) and the Granulocyte (G; CD11b/Ly6G) compartment by FACS analysis. The spleen was mostly comprised of HSC- and MPP-derived cells. CMP-derived cells were solely detectable 1 week after transplantation. CLP-derived cells were barely detectable. No data available for CMP- and CLP-derived cells 3, 8 or 16 weeks post transplantation.
Light-blue dots: HSC-derived clones, dark-blue dots: MPP-derived clones, green dots: CMP-derived clones and yellow dots: CLP-derived clones.

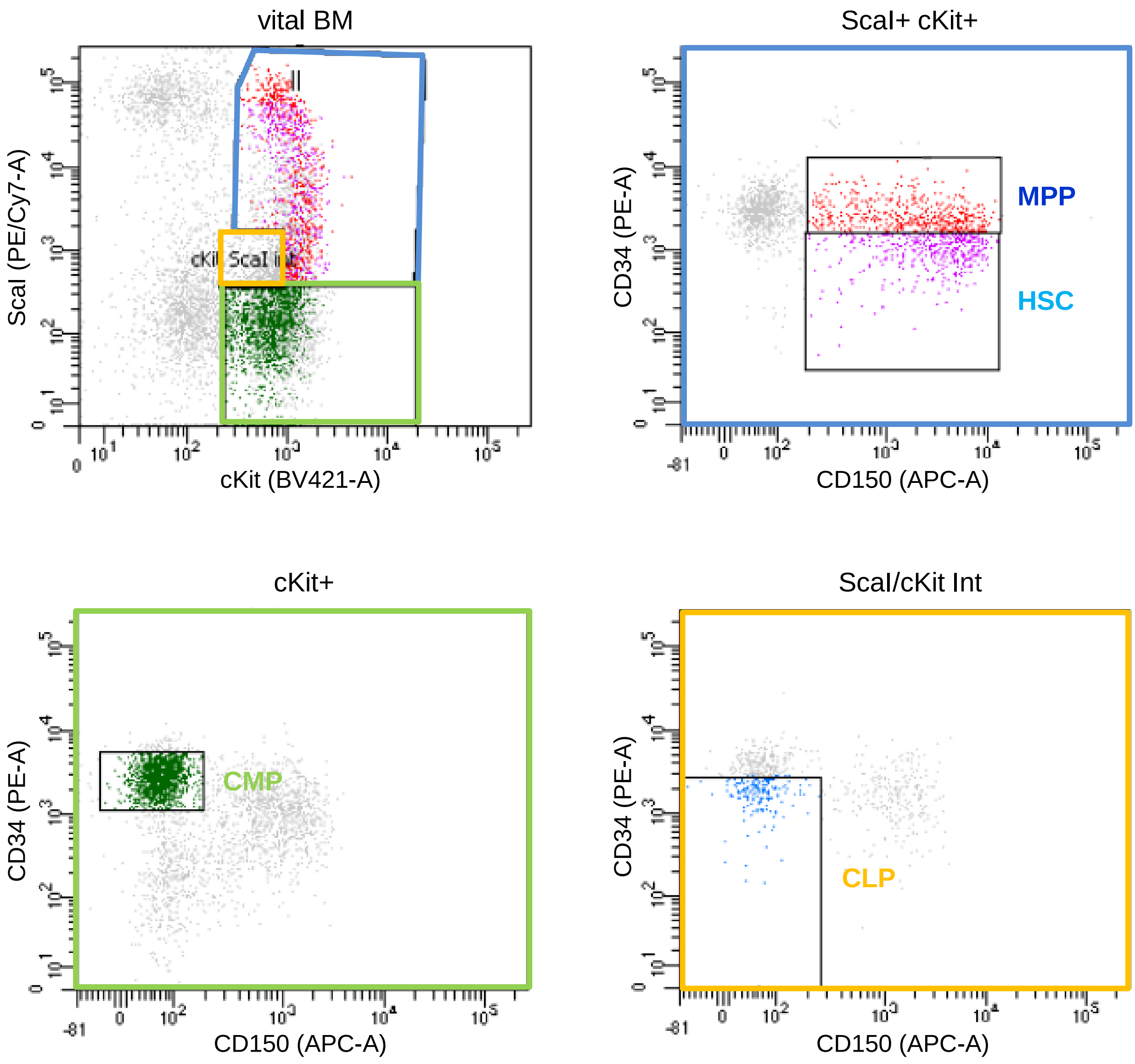

**Supplementary Figure 2: Sorting scheme of HSPC subpopulations based on their surface marker profile.** Bonemarrow cells were harvested from male donor animals and stained with the respective antibodies (Supplemental Table 2) against the surface markers ScaI, cKit, CD34 and CD150.
HSCs were defined as ScaI+ cKit+ CD150+ CD34-,
MPPs as ScaI+ cKit+ CD150+ CD34+,
CMPs as ScaI- cKit+ CD150- CD34+ and
CLPs as ScaI/ cKit Int CD150- CD34-.
Colors of the plots show parental gate in the vital BM plot and resemble color scheme of HSPC subpopulation used for visualization of the populations and their progeny.

**Supplementary Table 1: Antibodies used for fluorescence-activated cell sorting of HSPC populations and final analyses.**

| Name | Clone | Company | Application |
|---|---|---|---|
| anti CD16/32 | 93 | Biolegend | Sort of HSPC subpopulations/ Final Analyses |
| PE-Cy7 anti ScaI | E13-161.7 | Biolegend | Sort of HSPC subpopulations/ Final Analyses of BM populations |
| BV421 anti cKit | 2B8 | BD Biosciences | Sort of HSPC subpopulations |
| Pacific Blue anti cKit | 2B8 | Biolegend | Sort of HSPC subpopulations |
| APC anti CD150 | TC15-12F12.2 | Biolegend | Sort of HSPC subpopulations |
| PE anti CD34 | RAM34 | BD | Sort of HSPC subpopulations |
| APC-Cy7 anti cKit | 2B8 | Biolegend | Final Analyses of BM populations |
| PE anti CD150 | TC15-12F12.2 | Biolegend | Final Analyses of BM populations |
| APC anti lineage | diverse | BD (558074) | Final Analyses of BM populations |
| APC anti lineage | diverse | R&D Systems (FLC001A) | Final Analyses of BM populations |
| APC anti CD48 | HM48-1 | Biolegend | Final Analyses of BM populations |
| PE-Cy7 anti B220 | RA3-6B2 | Biolegend | Finaly analyses of mature blood cells in spleen |
| BV795 anti CD3e | 145-2C11 | BD | Finaly analyses of mature blood cells in spleen |
| PE anti Ly.6G | 1A8 | Biolegend | Finaly analyses of mature blood cells in spleen |
| APC anti CD11b | M1/70 | Biolegend | Finaly analyses of mature blood cells in spleen |

**Supplementary Table 2: Oligonucleotides used for Barcode library construction and analysis.** N positions indicate sites for random insertion of one of the four nucleotides during production, x indicates phosphothioate bond and BHQ1 stands for Black Hole Quencher 1.

| Internal name | Sequence (5‘-3‘) | Application |
|---|---|---|
| Poly-GFP-BC-fw | GGTGCATCTAGAACACTCTTTCCCTACACGACGCTCTTCCGATCTNNNACTNNCGANNCTTNNCGANNCTTNNGGANNCTANNACTNNCGANNCTTNNCGANNCTTNNGGANNCTANNACTNNCGANNCTCGAGGTGCACTATG | Oligo for GFP-BC library construction |
| Poly-Venus-BC-Fw | GGTGCATCTAGAACACTCTTTCCCTACACGACGCTCTTCCGATCTNNNCGANNAGANNCTTNNCGANNCTANNGGANNCTTNNCGANNAGANNCTTNNCGANNCTANNGGANNCTTNNCGANNAGANNCTCGAGGTGCACTATG | Oligo for Venus-BC library construction |
| Poly-Cerulean-BC-Fw | GGTGCATCTAGAACACTCTTTCCCTACACGACGCTCTTCCGATCTNNNCAGNNATCNNCTTNNCGANNGGANNCTANNCTTNNCAGNNATCNNCTTNNCGANNGGANNCTANNCTTNNCAGNNATCNNCTCGAGGTGCACTATG | Oligo for T-Sapphire-BC library construction |
| Poly-Cherry-BC-Fw | GGTGCATCTAGAACACTCTTTCCCTACACGACGCTCTTCCGATCTNNNCTANNCAGNNCTTNNCGANNCTANNCTTNNGGANNCTANNCAGNNCTTNNCGANNCTANNCTTNNGGANNCTANNCAGNNCTCGAGGTGCACTATG | Oligo for BFP-BC library construction |
| 32BC-Poly-fw | GGTGCATCTAGAACACTC | Generation of ds BC oligo |
| 32BC-Poly-rev | CATAGTGCACCTCGAG | Generation of ds BC oligo |
| Illu_P2 (43) | GTGACTGGAGTTCAGACGTGTGCTCTTCCGATCT | Bridging oligo for sequencing |
| Illu_MPLX35 (164) | CAAGCAGAAGACGGCATACGAGATTCTGAGGTGACTGGAGTTC | Index35 |
| Illu_MPLX36 (165) | CAAGCAGAAGACGGCATACGAGATCCTTGCGTGACTGGAGTTC | Index36 |
| Illu_MPLX37 (166) | CAAGCAGAAGACGGCATACGAGATTGGAGCGTGACTGGAGTTC | Index37 |
| Illu_MPLX38 (167) | CAAGCAGAAGACGGCATACGAGATTCGGGAGTGACTGGAGTTC | Index38 |
| Illu_MPLX39 (168) | CAAGCAGAAGACGGCATACGAGATAAACCT GTGACTGGAGTTC | Index39 |
| Illu_MPLX40 (169) | CAAGCAGAAGACGGCATACGAGAT CTCTAC GTGACTGGAGTTC | Index40 |
| Illu_MPLX41 (170) | CAAGCAGAAGACGGCATACGAGAT CGGCCT GTGACTGGAGTTC | Index41 |
| Illu_MPLX42 (171) | CAAGCAGAAGACGGCATACGAGATCCGGTGGTGACTGGAGTTC | Index42 |
| Illu_MPLX43 (172) | CAAGCAGAAGACGGCATACGAGATCAGCAGGTGACTGGAGTTC | Index43 |
| Illu_MPLX44 (173) | CAAGCAGAAGACGGCATACGAGATAAGTGCGTGACTGGAGTTC | Index44 |
| Illu_MPLX45 (174) | CAAGCAGAAGACGGCATACGAGATCAGGCCGTGACTGGAGTTC | Index45 |
| Illu_MPLX46 (175) | CAAGCAGAAGACGGCATACGAGATGGTAGAGTGACTGGAGTTC | Index46 |
| Illu_MPLX47 (176) | CAAGCAGAAGACGGCATACGAGATCCAGCAGTGACTGGAGTTC | Index47 |
| Illu_MPLX48 (177) | CAAGCAGAAGACGGCATACGAGATGCGCCAGTGACTGGAGTTC | Index48 |
| Illu_MPLX49 (178) | CAAGCAGAAGACGGCATACGAGATGGAACTGTGACTGGAGTTC | Index49 |
| Illu_MPLX50 (179) | CAAGCAGAAGACGGCATACGAGATGCGGACGTGACTGGAGTTC | Index50 |
| Illu_MPLX51 (180) | CAAGCAGAAGACGGCATACGAGATCGAAACGTGACTGGAGTTC | Index51 |
| Illu_MPLX52 (181) | CAAGCAGAAGACGGCATACGAGATCCACTCGTGACTGGAGTTC | Index52 |
| ILL_Dual_P5-01 | AATGATACGGCGACCACCGAGATCTACACAGCTTAGTACACTCTTTCCCTACACGACGCTCTTCCGATCxT | Dual Index01 |
| ILL_Dual_P5-02 | AATGATACGGCGACCACCGAGATCTACACGCTACTTGACACTCTTTCCCTACACGACGCTCTTCCGATCxT | Dual Index02 |
| ILL_Dual_P5-03 | AATGATACGGCGACCACCGAGATCTACACCTAGGCACACACTCTTTCCCTACACGACGCTCTTCCGATCxT | Dual Index03 |
| ILL_Dual_P5-04 | AATGATACGGCGACCACCGAGATCTACACTAGCAGCAACACTCTTTCCCTACACGACGCTCTTCCGATCxT | Dual Index04 |
| Y Chromosome Fw | ACAAGTTTTGGGACTGGTGACAA | ddPCR Y chromosome |
| Y Chromosome Rv | ACCACGGGACCACACCATAA | ddPCR Y chromosome |
| Y Chromosome-p | TGTCAAGCGCCCCATGAATGCA-BHQ1 | ddPCR Y chromosome |
| LeGOddBCi Fw | CTAGAAAAACATGGAGCAATCACAA | ddPCR illumina sequence |
| ddBCillu Rv | AGCGTCGTGTAGGGAAAGAGTG | ddPCR illumina sequence |
| ddBCIllu-p | CTGAATGATACGGCGACCACCGTCTAG | ddPCR illumina sequence |
| mEpo Chromosome Fw | GCAGGCGGGGTCGCTACTC | ddPCR mEpo |
| mEpo Chromosome Rv | CGCCTGTGCAGATCCGATAA | ddPCR mEpo |
| mEpo Chromosome-p | TTCTGAGGCGCCACTTTTGCAAGACC | ddPCR mEpo |

**Supplementary Table 3:**
Single cells were sorted based on their fluorescent protein expression. Cells were directly used for PCR to amplify the barcode sequence(s) using wpre-fw (5'-GAGGAGTTGTGGCCCGTTGT) and TA98 (5'-GGCTGTCAAACCTCCACTCT) for the first PCR and TA97 (5'-CATCGATACCGTCGACCTC) and TA99 (5'-TCGTTGGGAGTGAATTAGCC) for the second PCR. PCR-products were gelpurified and Sanger sequenced using TA99. The vector copy number (VCN) was determined as described in Bystrykh et al., 2016.

| Y-O | | | | | | O-O | | | | |
|---|---|---|---|---|---|---|---|---|---|---|
| HSCs | | | MPPs | | | HSCs | | | MPPs | |
| sample | VCN | | sample | VCN | | sample | VCN | | sample | VCN |
| 1 | 3,0 | | 1 | 3,0 | | 1 | 2,0 | | 1 | 2,0 |
| 2 | 3,5 | | 2 | 3,0 | | 2 | 2,0 | | 2 | 2,0 |
| 3 | 5,0 | | 3 | 5,5 | | 3 | 2,0 | | 3 | 4,0 |
| 4 | 2,0 | | 4 | 3,5 | | 4 | 1,0 | | 4 | 3,0 |
| 5 | 2,0 | | 5 | 4,0 | | 5 | 1,0 | | 5 | 5,5 |
| 6 | 3,0 | | 6 | 4,0 | | 6 | 1,0 | | 6 | 2,0 |
| 7 | 2,0 | | 7 | 2,0 | | 7 | 4,5 | | 7 | 3,0 |
| 8 | 3,0 | | 8 | 2,0 | | 8 | 1,0 | | 8 | 1,0 |
| 9 | 1,0 | | 9 | 2,5 | | 9 | 1 | | 9 | 2,0 |
| 10 | 3,0 | | 10 | 2,5 | | 10 | 1 | | 10 | 2,0 |
| 11 | 2,0 | | 11 | 3,0 | | 11 | 2 | | 11 | 5,0 |
| 12 | 4,0 | | 12 | 3,5 | | 12 | 2 | | 12 | 2,0 |
| 13 | 4,0 | | 13 | 3,0 | | 13 | 2 | | 13 | 1,0 |
| 14 | 4,0 | | 14 | 5,5 | | 14 | 2 | | 14 | 1,0 |
| 15 | 1,0 | | 15 | 1,0 | | 15 | 3 | | 15 | 1,0 |
| 16 | 6,0 | | 16 | 5,0 | | 16 | 2 | | 16 | 1,00 |
| 17 | 1,0 | | 17 | 4,0 | | 17 | 4 | | 17 | 2,00 |
| 18 | 4,0 | | 18 | 6,0 | | | | | 18 | 2,00 |
| 19 | | | 19 | 1,0 | | | | | 19 | 3,00 |
| 20 | | | 20 | 1,0 | | | | | 20 | 1,00 |
| 21 | | | 21 | 3,0 | | | | | | |
| mean | 3,0 | | mean | 3,2 | | mean | 2,0 | | mean | 2,3 |
